# A non-canonical-PPARγ/RXRα-binding sequence regulates leptin expression in response to changes in adipose tissue mass

**DOI:** 10.1101/325480

**Authors:** Yinxin Zhang, Olof Stefan Dallner, Tomoyoshi Nakadai, Gulya Fayzikhodjaeva, Yi-Hsueh Lu, Mitchell A. Lazar, Robert G. Roeder, Jeffrey M. Friedman

## Abstract

Leptin expression decreases after fat loss and is increased when obesity develops and its proper quantitative regulation is essential for the homeostatic control of fat mass. We previously reported that a distant leptin enhancer (LE1), 16kb upstream from the transcription start site (TSS), confers fat-specific expression in a BAC transgenic reporter mouse (BACTG). However this and the other elements that we identified do not account for the quantitative changes in leptin expression that accompany alterations of adipose mass. In this report, we used ATAC-seq to identify a 17bp non-canonical-PPARγ/RXRα-binding site leptin regulatory element 1 (LepRE1) within LE1, and show that it is necessary for the fat-regulated quantitative control of reporter (luciferase) expression. While BACTG reporter mice with mutations in this sequence still show fat-specific expression, luciferase is no longer decreased after food restriction and weight loss. Similarly the increased expression of leptin reporter associated with obesity in *ob/ob* mice is impaired. A functionally analogous LepRE1 site is also found in a second, redundant DNA regulatory element 13kb downstream of the TSS. These data uncouple the mechanisms conferring qualitative and quantitative expression of the leptin gene and further suggest that factor(s) that bind to LepRE1 quantitatively control leptin expression and might be components of a lipid sensing system in adipocytes.

**Significance:** Leptin gene expression is highly correlated with the lipid content of individual fat cells suggesting that it is regulated by a "fat sensing" signal transduction pathway. This study is thus analogous to studies that led to the identification of a cholesterol-sensing pathway by studying the regulation of the LDL receptor gene by intracellular cholesterol. Several lines of investigation have suggested that, in addition to adipocytes, liver, neurons and other cell types can also sense changes in lipid content though the molecular mechanisms are unknown. The data here provide a critical first step toward elucidating the components of this system, which would be of great importance. These studies also identify a previously underappreciated role of PPARγ/RXRα complex to regulate leptin expression.

## Introduction

Leptin is an adipocyte hormone that maintains homeostatic control of adipose tissue mass and functions as an afferent signal in a negative feedback loop. Leptin deficiency leads to extreme obesity both in mouse and human while leptin treatment of wild type mice reduces fat mass (1–7). Weight gain leads to an increased leptin level that in turn inhibit food intake and return adipose tissue mass to the starting point. Similarly, weight loss decreases the leptin level, leading to increased food intake and increased fat mass. Thus the quantitative changes in leptin expression associated with changes in nutritional state are critical for the proper functioning of this system. Consistent with this, the levels of leptin RNA and plasma leptin are highly correlated with adipocyte cell size and cellular lipid content (8–10). This has suggested that, analogous to a cholesterol sensing system in liver and other cell types (11, 12), there might be a lipid sensing mechanism (or another mechanism such as sensing of cell size) in adipocytes that adjusts the level of leptin gene expression to changes in the level of lipid stores. However, neither the molecular mechanisms controlling the change in leptin expression levels nor the elements of this putative signal transduction pathway are known (13).

To address this question, we initiated efforts to define cis elements and trans factors that control the quantitative expression of the leptin gene. Defining DNA sequence binding sites has been crucial for identifying numerous transcription factors, including SP1(14) and NF-kB(15). This approach is also analogous to that used to identify the aforementioned cholesterol sensing system. In that case, studies of LDL receptor expression revealed that the levels of cholesterol in the ER membrane regulate cleavage and nuclear transport of the SREBP transcription factor, in turn controlling the expression of genes that regulate cholesterol metabolism. However, analogous studies of leptin require that expression be monitored in vivo because cultured adipocytes, which have a very low lipid content, express an approximately thousand-fold lower level of leptin RNA than do fat cells in vivo (16). Previous efforts to map cis elements controlling leptin expression have thus used Bacterial Artificial Chromosomes transgenic reporter animals (BACTGs) with luciferase inserted at the translation start site of the leptin gene (17). Through a comprehensive deletion analysis, we previously found that a BACTG encompassing 31kb (−22 to +8.8 kb) of the leptin locus showed fat-specific luciferase expression (18). Similar to the endogenous gene, luciferase expression was reduced after 48h of food deprivation while its expression was increased after crossing of the reporter mice to ob/ob mice that express very high levels of leptin RNA. We also found that a 3’ BACTG extending from −762bp to +18kb was able to drive reporter expression in a manner similar to the 5’ 31kb (−22 to +8.8 kb) BACTG, while a BACTG extending from −762bp to 8.8kb lost fat-specific reporter expression. Subsequent studies have identified two redundant elements that can independently confer cell-specific expression of leptin in adipocytes; LE1 is localized between −16.5 and −16.1 kb upstream of the leptin TSS while LE2 is located between + 13.6 and +13.9 kb downstream of the leptin TSS(19). However, because deletion of these elements ablates fat-specific expression altogether, it is not possible to assess the role of these specific sequences to quantitatively regulate leptin expression. Similarly, while other factors including C/EBPα and SP1(20, 21), FOSL2 (22) and NFY (18), have been reported to play a role in leptin expression, none have been shown to quantitatively regulate this gene.

In this report, we employed the Assay for Transposase Accessible Chromatin with high - throughput sequencing (ATAC-seq) (23) as part of an unbiased screen to identify sites of transcription factor binding in inguinal white adipose tissue (iWAT) from fed, food restricted (48h), and ob/ob mice. Deep sequencing revealed a highly conserved non-canonical footprint for a PPARγ/RXRα binding site within LE1 (referred to hereafter as LepRE1 or Leptin Regulatory Element 1) in adipose tissue nuclei derived from ob/ob mice, but not from adipose tissue of wild type mice. Point mutations in this regulatory sequence in the 5’ reporter BACTG (−22 to 8.8kb) abrogated quantitative regulation of luciferase in adipose tissue from fasted and obese mice. A functionally equivalent PPARγ/RXRα-binding site was also found within LE2, potentially explaining the functional redundancy of these elements. Purified PPARγ/RXRα binds weakly to these sequences and both gain of function and loss of function mutations within the core sequence affect the regulated expression of the reporter. In addition, mutations in the adjacent PPARγ extension site also disrupt the proper quantitative control of reporter expression. These data suggest a model in which the quantitative regulation of leptin expression depends on the stabilization of PPARγ/RXRα binding to an otherwise weak binding site by an accessory factor binding to the adjacent extension site.

## Results

**A Fat-Regulated Footprint LepRE1 is a Non-Canonical-PPARγ/RXRα-Binding Site.** In order to find potential leptin regulatory elements that respond to changes in fat mass, we isolated nuclei from inguinal white adipose tissue (iWAT) of fasted, fed and ob/ob mice and generated genome-wide footprints using ATAC-seq. In this method, isolated nuclei are incubated with a hyperactive transposase loaded with adaptor sequences so that DNA sequences from open regions of chromatin (i.e., DNase hypersensitive regions) can be PCR amplified using the adaptor sequences as primers. Deep sequencing of the PCR products yields a genome wide inventory of sequences from accessible regions of chromatin (23). Furthermore, analysis of the heights of the peaks of the DNA sequence reads reveals DNA footprints indicative of protein binding. Using this method, we identified six peaks within −22 to +8.8 kb of the leptin gene that showed a 3-fold or higher signal in adipose tissue nuclei from ob/ob mice compared to nuclei from fed and fasted mice (Fig 1A). These six peaks included the aforementioned LE1 between −16.5 and −16.1 kb of the TSS, the proximal promoter around the TSS, and four regions within the first intron. The differences in access of the transposase to the proximal promoter and the transcribed regions in the first exon are consistent with the higher level of expression of this gene in ob vs. wild type adipose tissue.

**Fig 1.**
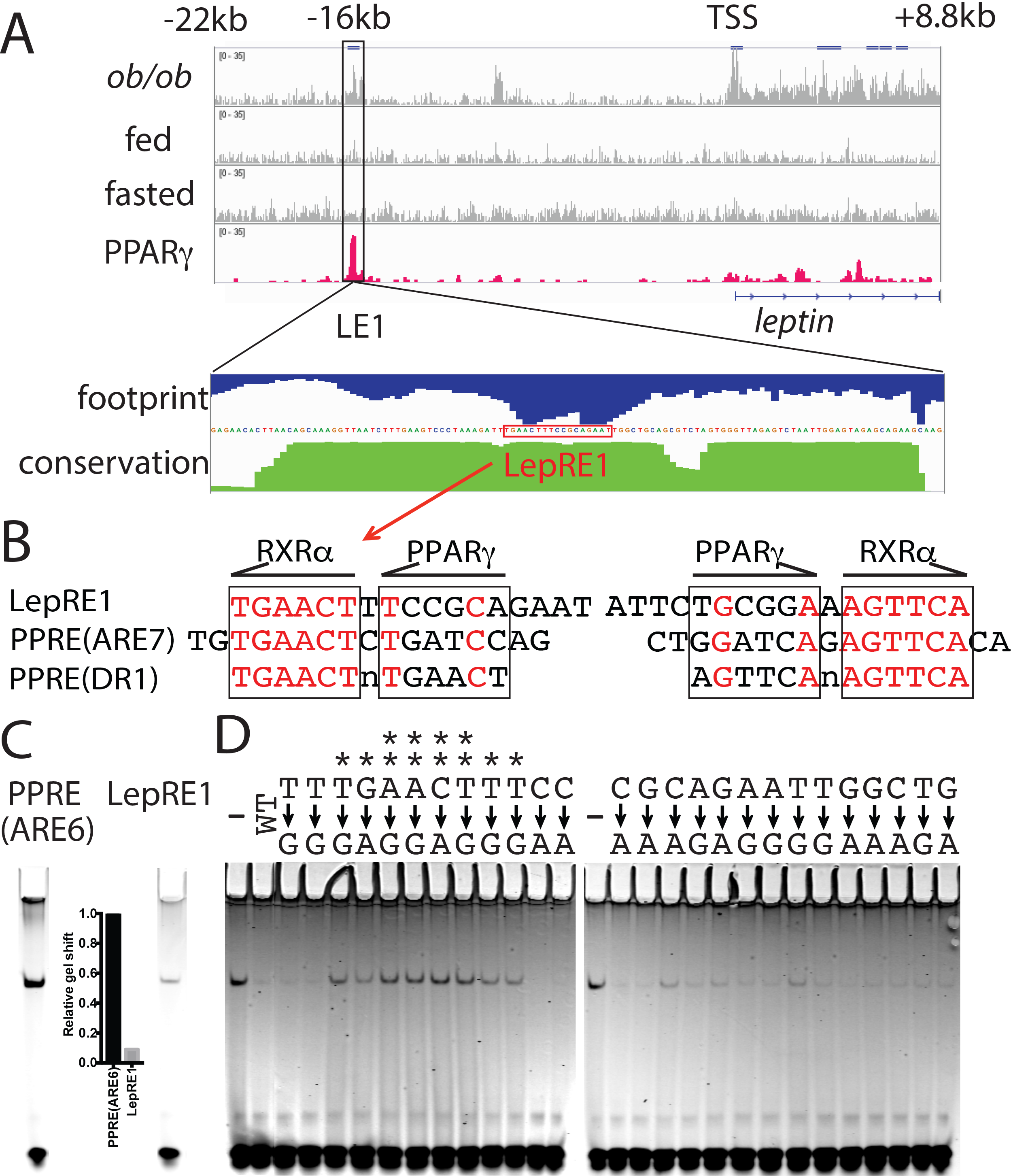
A Fat-Regulated Footprint LepRE1 is a Non-Canonical-PPARγ/RXRα-Binding Site. A. ATAC-seq was performed using nuclei from the inguinal fat of ob/ob, fed and fasted B6 mice, and compared to PPAR γ ChIP-seq in the inguinal fat from B6 mice (33, 34). Data for sequences between −22kb to +8.8kb sequences of the leptin gene locus are shown. Regions with a 3 fold change from ob/ob mice vs. fasting are highlighted. The −16kb region of leptin gene is enlarged to show the calculated footprint score (Blue) and conservation score within mammals (Green). The LepRE1 sequence is denoted by red box. B. Sequence alignment of LepRE1, PPRE(ARE7) and PPRE(DR1). The LepRE1 sequence denoted by red box in A is shown on the left. The nucleotides that are conserved between the PPREs and LepRE1 are in red. PPARγ and RXRα binding sequences are boxed based on solved crystal structure(41). C. EMSA of purified PPARγ/RXRα with IRDye 700 labeled PPRE(ARE6) and LepRE1 oligos. The shifted bands were quantified with ImageStudioLite and normalized to PPRE(ARE6). D. EMSA competition studies of excess unlabeled wild type LepRE1 oligos or oligos with scanned point mutations with IRDye 700 labeled wild type LepRE1 in the presence of purified PPARγ/RXRα.

We thus focused on the LE1 sequence at ~ −16kb because it is not transcribed and also because a 400 bp deletion of LE1 in a BAGTG (−22 to 8.8kb) abolishes fat specific expression of a luciferase reporter. Within LE1 there is a 101bp segment (mm9, chr6: 28993757-28993857) that is almost identical among twenty placental mammal species, which is consistent with the finding that fat-specific expression of leptin is only evident in mammals (24). Within this 101 bp region of LE1, there is a small 17bp footprint in ob/ob but not in wild type adipose tissue nuclei. The footprint showed two apparent peaks, suggesting that two (or more) proteins bound there. Hereafter, we refer to this footprinted sequence as Leptin Regulator Element 1 (LepRE1) (Fig 1A).

We noticed that 6 consecutive base pairs (out of 13) of the footprinted sequence of LepRE1 were identical to the RXRα-binding sequence (referred to as the conserved DR1 half-site) of the Fabp4/aP2 gene enhancer. This binding site is referred to as the Peroxisome Proliferator Response Element/Adipocyte Regulatory Factor Response Element 7 (also known as ARE7) and this binding site is composed of a direct repeat of two DR1 half-sites with 1-nucleotide spacer (25, 26)(Fig 1B). PPARγ, a member of nuclear hormone receptor superfamily, is the master regulator of adipogenesis (27, 28). This transcription factor binds as a heterodimer with RXRα, another nuclear receptor, to a Direct Repeat 1(DR1) sequence (29). However in LepRE1, the next 6 bp sequence, 3’ to the conserved DR1 (i.e; the second half site) had only very limited homology to DR1 as would typically be found in a canonical PPARγ-binding sequence or PPRE (Peroxisome Proliferator Response Element).

Thus the footprint we found in the −16kb upstream region of the leptin gene is not a canonical PPRE (DR1) in that a single DR1 half-site was followed by a non-homologous sequence (Fig 1B and Fig S1A). Indeed, because of this, none of the current algorithms that identify DNA binding sites identified LepRE1 as a PPARγ/RXRα-binding sequence at all. However, as shown below, this sequence can bind to a PPARγ/RXRα heterodimer, albeit more weakly than does a canonical binding site.

We confirmed that the PPARγ/RXRα protein complex could bind to this sequence using an EMSA assay in which purified PPARγ and RXRα were incubated with LepRE1 oligonucleotides labeled with the infrared dye (IRDye) 700. FLAG-tagged mouse PPARγ2 and human RXRα proteins were purified from Baculovirus infected Sf9 cells. We found that the purified protein bound to this sequence with an affinity one tenth of that of a canonical DR1 PPRE. In these studies, we used the ARE6 binding site rather than the aforementioned ARE7 binding site because more extensive mutagenesis has been performed for this sequence (25)(Fig 1C). ARE6 and 7 share six of thirteen core DR1 nucleotides and both provide high affinity sites for PPARγ binding to the Fabp4/aP2 enhancer (26). This gel shift was abolished after co-incubation with an excess of an unlabeled wild type DNA fragments, while unlabeled DNA fragments with point mutations in the RXRα-binding sequence -- in particular the AACT/AGTT part of the conserved DR1 half-site of LepRE1 -- no longer competed with the labeled probe in the gel shift (Fig 1D). In contrast, unlabeled DNA fragments with point mutations in the sequences 3’ to the conserved DR1 half-site, the PPARγ-binding sequence (TCCGCA/TGCGGA), as well as mutations in the extension sequence for PPARγ (Fig 1D) competed for binding in a similar manner to wild type oligonucleotides. The extension sequence is adjacent to the non-conserved half site with the sequence GAAT/ATTC. It previously has been suggested that this site binds other transcription factors, the identity of which have not been determined (30). These results show that the RXRα-binding portion of LepRE1 is required for the (weaker) binding of PPARγ/RXRα to this sequence (Fig 1D). These in vitro EMSA findings are also consistent with data from ChIP-seq analyses identifying sites of PPARγ binding (31–34)(Fig 1A and Fig S1B). These data confirm that PPARγ/RXRα weakly binds to LepRE1 in adipocyte nuclei in vivo via interactions with a non-canonical PPARγ binding site.

**A 3’ Fat-Regulated Footprint, LepRE2, is Homologous to LepRE1**. LE1, located between −16.5 to −16.1kb, and LE2, located between +13.6 to +13.9 kb, are redundant elements that can independently confer qualitative and quantitative expression of luciferase in leptin-reporter mice -- raising the possibility that functionally similar elements at both sites can regulate the level of leptin gene expression(18, 19). In previous studies, deletion of LE2 in a BAGTG extending between −0.762 to +18 kb ablated fat specific expression of a luciferase reporter. We thus used ATAC seq to identify open regions of chromatin within −0.762 to +18 kb region of the leptin gene and found ten hypersensitive peaks with a three-fold difference between ob/ob and fasted mice. All but one of these ten peaks were in transcribed regions that included the proximal promoter around the TSS and four regions within the first intron (referred to previously), as well as one region within the second intron, two regions within the third exon, one region around the transcription termination site, and one region downstream the transcription termination site.

The only site from a non-transcribed region was found immediately after the transcription termination site, and showed a highly significant peak in ob/ob adipocyte nuclei in a 66bp segment of LE2 (mm9, chr6: 29023967-29024032). As was also the case for LepRE1, this sequence was conserved among twenty placental mammal species (Fig 2A). The ATAC-seq data identified a small footprint within this 66bp segment referred to as LepRE2 (Fig 2A). LepRE2 was identical to LepRE1 at ten of seventeen (59%) nucleotides (Fig 2B). Among the ten identical nucleotides between LepRE1 and LepRE2, four (ATTC/GAAT) were in the 5’ extension sites of PPARγ while five (GAAAG/CTTTC of the other six were in the middle of the PPARγ/RXRα binding site (Fig 2B). Although the sequence LepRE2 did not contain a single conserved DR1 half-site sequence as in LepRE1, LepRE2 had the same DNA sequence (AAAGG/CCTTT) as a central portion of a consensus DR1 motif as shown in the analyses of genome wide PPARγ ChIP-seq and RXRα ChIP-seq data (31, 32) (Fig S1A). Like LepRE1, current algorithms that identify DNA binding sites did not designate this LepRE2 sequence as a PPARγ/RXRα-binding sequence. Purified PPARγ/RXRα complex bound to this sequence as shown by EMSA (Fig 2C). However, similar to LepRE1, the binding affinity of purified PPARγ/RXRα complex to an IRDye 700-labeled LepRE2 oligonucleotide was also weak with an approximately five-fold lower signal compared to that observed for binding to a typical PPRE (ARE6). The binding site of the PPARγ/RXRα complex was further refined by testing the ability of excess unlabeled wild type oligonucleotides and oligonucleotides with mutations in each of the positions in LepRE2 to compete with the labeled oligonucleotide binding to PPARγ/RXRα (Fig 2D). Oligonucleotides with point mutations in the central part of DR1 ChIP-seq motif (AAAGG/CCTTT) no longer competed with the gel shift band, while mutations in other segments that included the extension sequence showed a similar ability as wild type oligonucleotides to compete for binding. Similar to LE1/LepRE1, the LE2/LepRE2 peak was also identified in several published PPARγ ChIP-seq datasets(33, 34), indicating binding of this transcription factor to the leptin gene in adipocytes in vivo (Fig 2A and Fig S1B).

**Fig 2.**
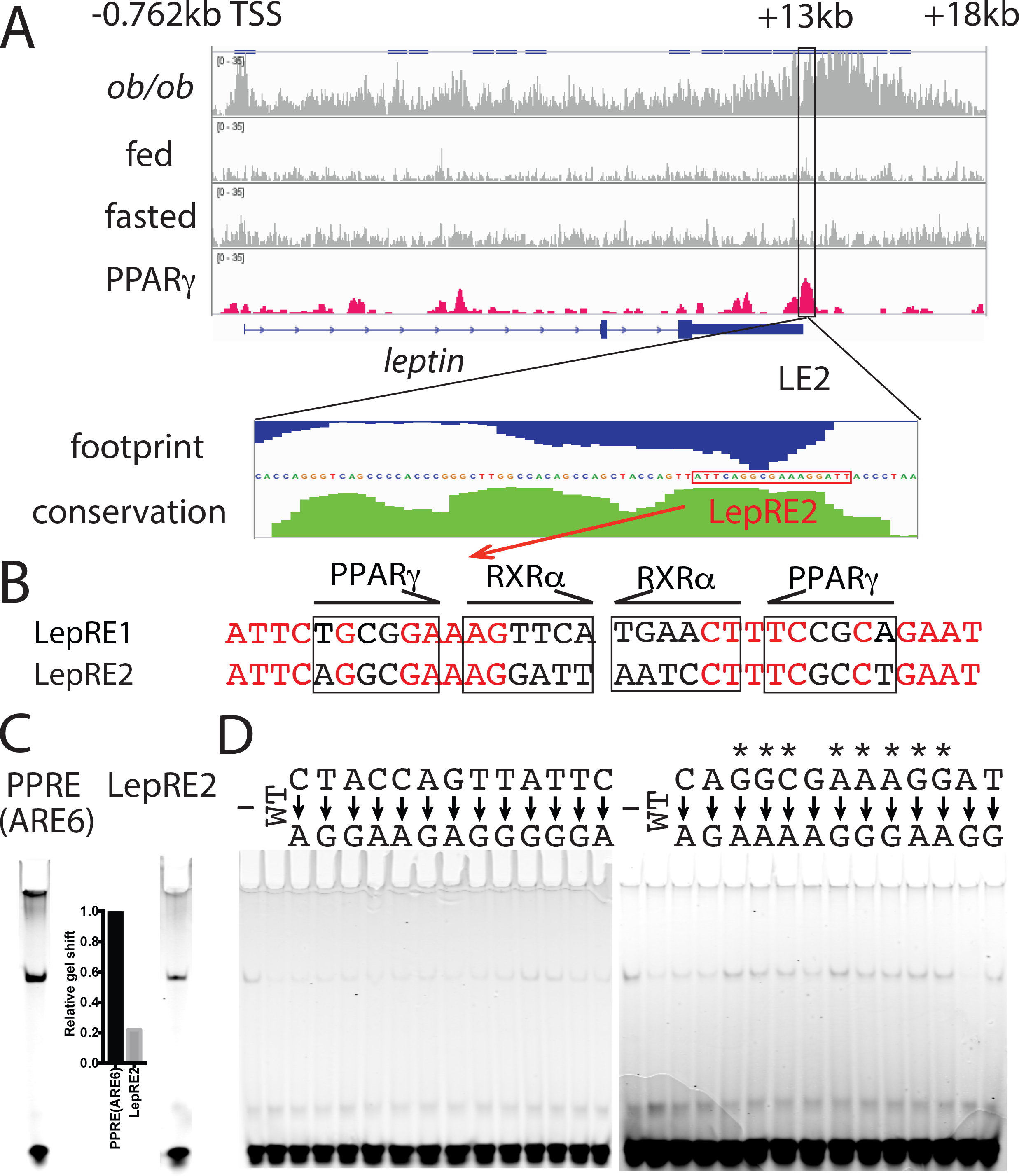
A 3’ Fat-Regulated Footprint, LepRE2, is Homologous to LepRE1. A. ATAC-seq results of the inguinal fat from ob/ob, fed and fasted B6 mice, and PPARγ ChIP-seq in the inguinal fat from fed B6 mice (33, 34) between −0.762kb and +18kb sequences of the leptin locus are shown. Regions with a 3 fold change in nuclei from ob/ob vs. fasted mice are highlighted. The +13kb region of the leptin gene is enlarged to show the calculated footprint score (Blue) and conservation score within mammals (Green). The LepRE2 sequence is denoted by a red box. B. Sequence comparison of LepRE1 and LepRE2. The LepRE2 sequence denoted by red box in A is shown on the left. The conserved nucleotides are red. PPARγ and RXRα binding sequences are boxed based on solved crystal structure(41). C. EMSA of purified PPARγ/RXRα with IRDye 700 labeled PPRE(ARE6) and LepRE2 oligos. The shifted bands were quantified with ImageStudioLite and normalized to PPRE(ARE6). D. EMSA competition studies of excess unlabeled wild type LepRE2 and oligos with point mutations using IRDye 700 labeled wild type LepRE2 in the presence of purified PPARγ/RXRα.

**Fat Specific Expression in Reporter Mice with LepRE1 Mutations.** Previous studies have shown that PPARγ, at least by itself, is not sufficient to induce a high level of leptin expression. For example, leptin expression is extremely low in cultured adipocytes and in brown fat despite the fact that they express high levels of this transcription factor (16). In order to confirm that LepRE1 and LepRE2 are functional PPREs, we co-transfected PPARγ and RXRα into HEK293T cells expressing a luciferase reporter construct. We found that PPARγ and RXRα can indeed activate the LE1 and LE2 enhancers upstream of a luciferase reporter with a ~10 fold induction relative to control experiments without co-transfection of PPARγ and RXRα (Fig 3A). Interestingly, treatment with the RXR ligand 9-cis-Retinoic acid decreases the LE1-driven and LE2-driven luciferase activity. The PPARγ ligand rosiglitazone also decreases reporter expression though to a lesser extent than 9-cis-Retinoic acid (Fig 3A). These results are consistent with the observation that thiazolidinediones have been reported to inhibit leptin expression (35–37).

**Fig 3.**
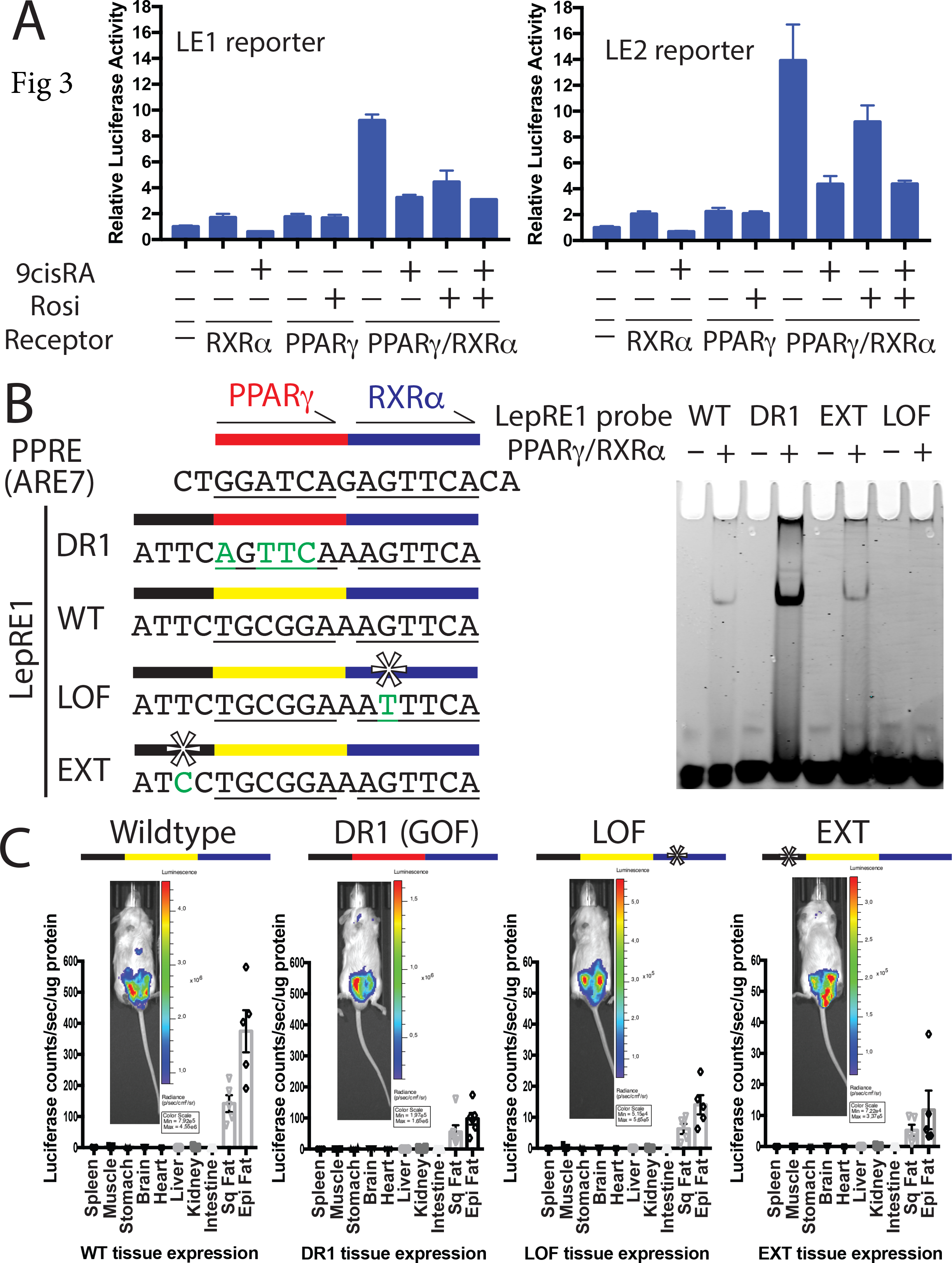
Fat Specific Expression in Reporter Mice with LepRE1 Mutations. A. PPARγ/RXRα complex activates both LE1 and LE2 enhancers in transfected HEK293T cells in dual luciferase reporter assay. B. A series of point mutations were introduced into LepRE1 for functional studies in vitro and in vivo. Sequence alignment of PPRE(ARE7), a LepRE1 DR1 gain of function mutation (GOF), wild type LepRE1, a LepRE1 LOF loss of function mutation, and a mutant in the LepRE1 extension site, EXT. The mutated nucleotides are green and an asterisk marks the sites of the mutations. A red line above the sequence indicates canonical PPARγ binding region, while a yellow line shows non-canonical PPARγ binding. The blue line shows RXRα binding. The black line represents the 5’ extension sequence. EMSA results of LepRE1 WT, DR1, LOF and EXT with purified PPARγ/RXRα are shown (right). C. These point mutations were then introduced into leptin luciferase reporter mice that extended between −22kb and +8.8kb of the leptin locus. Luciferase expression is shown for BAC Transgenic mice with the wild type, DR1, LOF and EXT LepRE1 sequences. 10 tissues were dissected, and processed for protein quantification and luciferase measurement. A representative IVIS image of mouse is also shown for each transgenic line.

We thus investigated whether this non-canonical PPARγ/RXRα binding site is required for proper qualitative and quantitative expression of leptin by making point mutations in LepRE1 in the reporter BACTG that extends from −22 to +8.8 kb. In this BACTG, luciferase is inserted by homologous recombination into the translation start site. Three LepRE1 mutants were characterized: i. a gain of function (GOF) PPARγ/RXRα-binding mutant LepRE1 DR1, in which a canonical DR1 half-site has replaced the second non-canonical 3’ binding site; ii. a loss-of-function PPARγ/RXRα-binding mutant LepRE1 (LOF) in which a point mutation was introduced into the highly conserved DR1 half-site sequence i.e; the RXRα-binding region that is required for binding (Fig 1D); and iii. an extension mutant LepRE1 EXT in which a point mutation was introduced in the 5’-flanking sequence upstream of the core DR1. As mentioned, previous studies suggest the 5’ extension site of PPARγ may be a target of other binding proteins (30). EMSA assays confirmed that the DR1 gain-of-function mutant bound to purified PPARγ/RXRα more strongly than did the wild type sequence, that the LOF mutant failed to bind, and that the EXT mutant bound with similar affinity to the wild type oligonucleotide (Fig 3B).

We next made multiple BACTGs reporter lines for each of the three different LepRE1 mutants. All of the lines for the gain-of-function DR1, loss-of-function (LOF) and extension (EXT) mutations expressed leptin luciferase reporter exclusively in adipose tissue, albeit with lower baseline levels of expression vs. the wild type reporter mice. In temporal analyses (Fig S2), we also noticed a progressive diminution in luciferase expression between three and eight weeks age with the relative level of luciferase expression in the LepRE1 DR1 reporter mice decreasing from 120% of the wild type BACTG to 20~30% that of the control construct by 8 weeks of age, after which time the expression level was stable. At baseline, the level of luciferase expression in the LepRE1 LOF reporter mice was fat-specific but was expressed at a considerably lower level of ~ 4-8% that of the wild type construct. Finally luciferase expression in the LepRE1 EXT reporter mice was ~15% that of the wild type reporter and decreased to a level ~4% that of the wild type reporter mice between 3 and 8 weeks of age (Fig S2). These data show that the newly identified non-canonical PPARγ/RXRα-binding site is necessary for normal levels of expression of the leptin gene. As mentioned, in all cases luciferase was expressed exclusively in adipose tissue and after eight weeks the level of luciferase expression was stable in all three lines. Thus, all subsequent experiments were performed in mice that were nine to thirteen weeks of age (Fig 3C). We next evaluated the impact of the PPARγ/RXRα-binding site mutations on the level of reporter expression after a period of food restriction or in obese animals.

**Dysregulation of Leptin Reporter Expression After Weight Loss and Weight Gain.** Weight loss after a period of food restriction is associated with a decrease in leptin gene expression and leptin plasma level. After two days of food restriction, similar to expression of the endogenous gene, expression of the leptin luciferase reporter in the 5’ WT BACTGs (−22 to +8.8 kb) decreased 3.3 fold (3.1x10^8^ p/s units of total flux) relative to the average expression level before fasting (Fig 4A). In contrast, the level of luciferase expressed from the gain of function mutation (with the stronger-PPAR γ/RXRα-binding site, LepRE1 DR1) did not decrease after two days of food restriction. Similarly, the reporter animals with a point mutation in the 5’ extension site (LepRE1 EXT) also showed a stable level of luciferase expression after a fast. Importantly, as shown previously (see Fig 3B) this mutation in the extension sequence did not alter the binding of PPARγ/RXRα to LepRE1. Finally, the reporter animals with a loss of function of PPARγ/RXRα-binding (LepRE1 LOF) showed a 1.6 fold decrease of luciferase expression after fasting (1.1x10^7^ p/s (total flux)), and the magnitude of this decrease was 27.7 fold lower than that seen in WT mice after two days of food restriction (Fig 4A and Fig 4B). The body weight and fat mass of each of the groups (11 to 13 weeks old) was the same showing that the altered reporter expression was not a result of differences in adipose tissue mass (Fig 4C).

**Fig 4.**
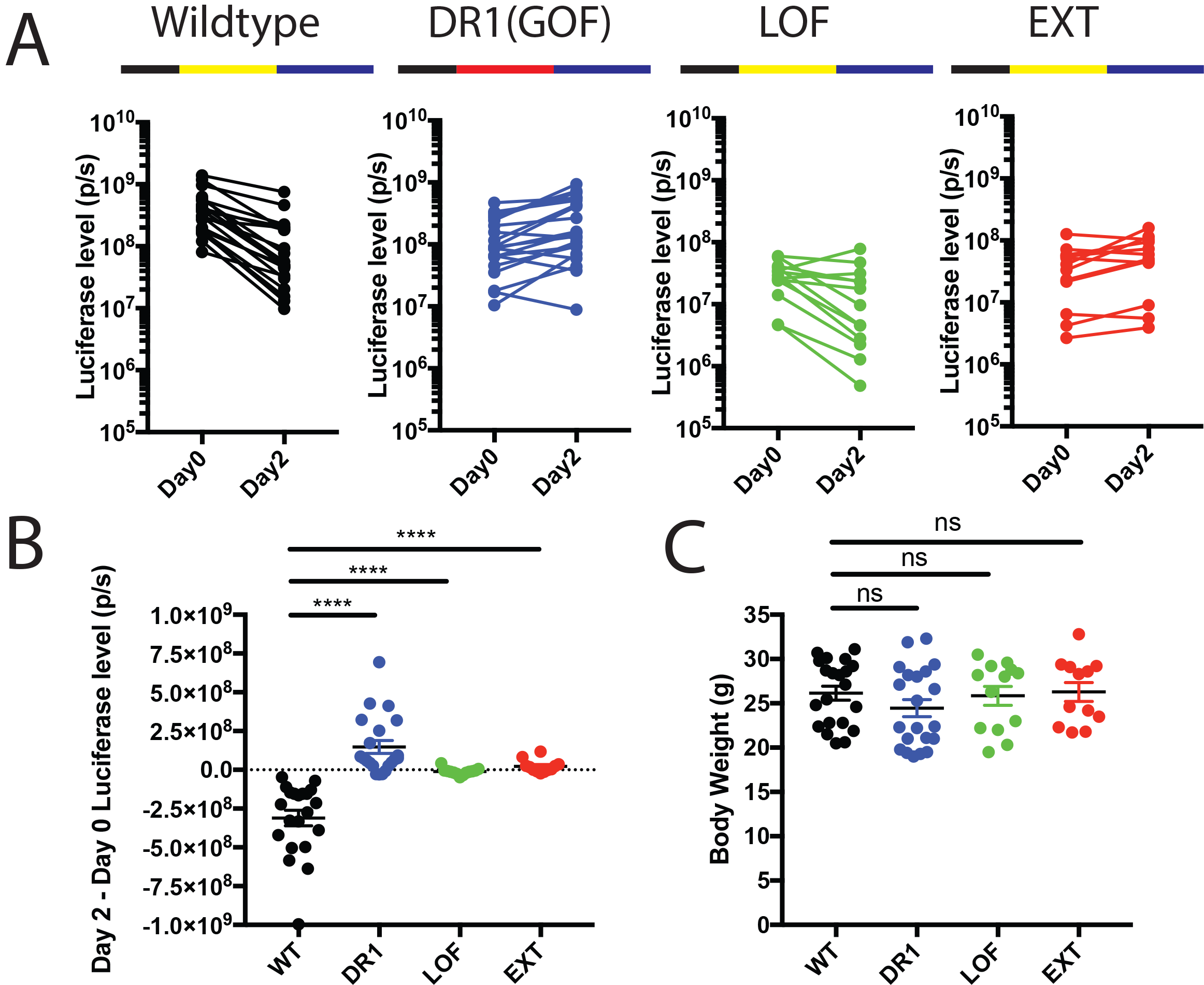
Dysregulation of Leptin Reporter Expression After Weight Loss. A. Individual whole body luciferase levels before fasting (Day 0) and after 2 days of fasting (Day 2) for the wild type leptin BAC (−22 to +8.8kb) and the DR1 (GOF), LOF and EXT mutants are shown. The mice were between 11 week old to 13 week old. B. The difference between Day 2 and Day 0 whole body luciferase level (Day 2-Day 0) for individual mice is shown. C. Body weight for corresponding individual mice on Day 0. For A-C, WT n=21, DR1 n=21, LOF n=13, EXT n=12.

We next analyzed the effect of these mutations on reporter expression in obese animals by mating the aforementioned reporter lines to ob/ob mice. ob/ob mice carry a mutation in the leptin coding sequence and show a dramatic compensatory increase in the level of expression of leptin RNA. As above, no significant differences in body weight were observed in the mutant reporter animals vs. mice carrying the wild type reporter (Fig 5A). Nine-week-old ob/ob mice had an average body weight 42g, which is significantly higher than the 21g average body weight in nine-week-old ob/+ mice. As previously reported, the level of the luciferase reporter was 9.6 fold higher (7.6×10^9^ p/s in total flux) in ob/ob transgenic animals expressing the wild type reporter relative to the level of reporter expression in non-obese ob/+ mice.

**Fig 5.**
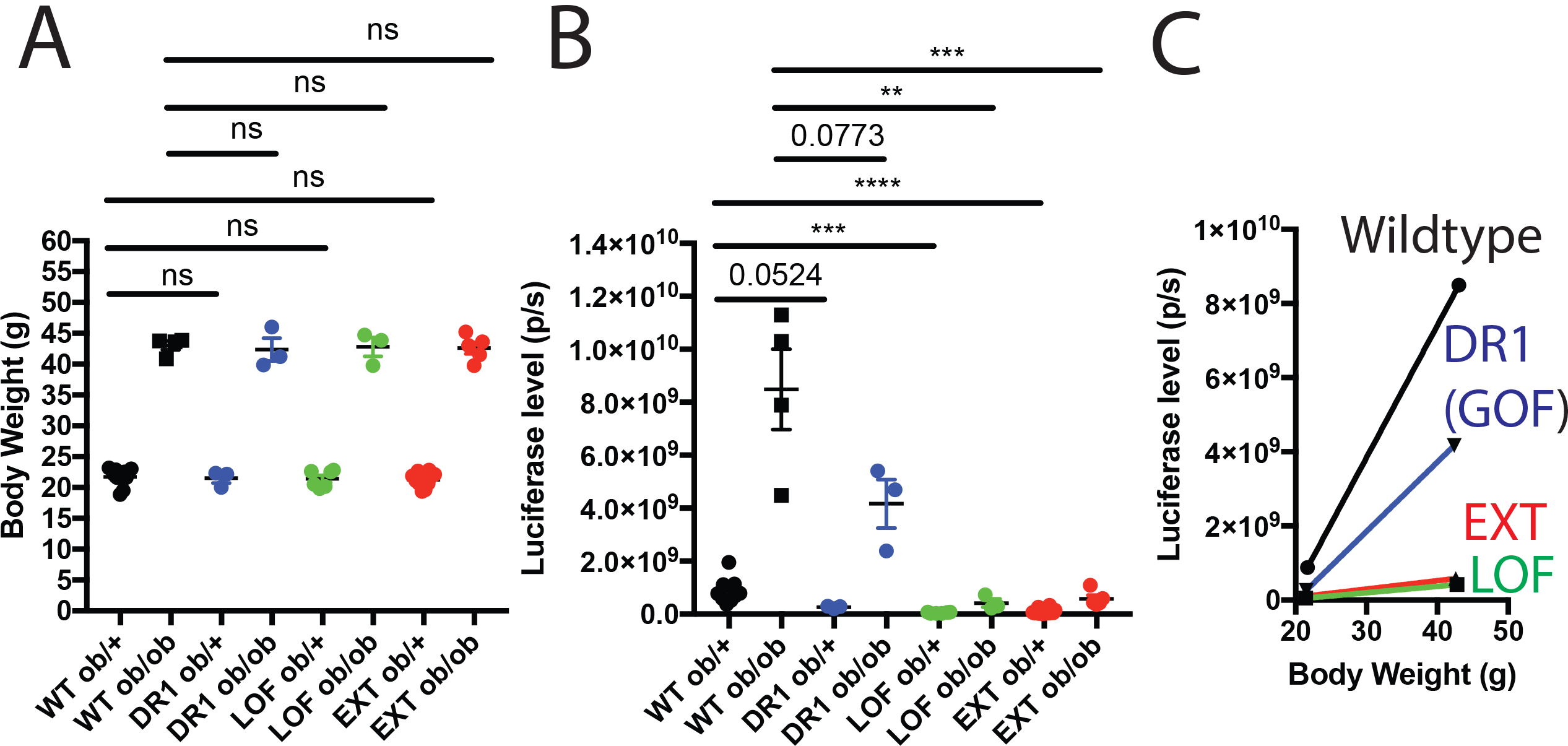
Dysregulation of Leptin Reporter Expression After Weight Gain. A. Body weight for individual 9-week ob/+ or ob/ob BAC Transgenic mice. B. The corresponding whole body luciferase level for individual 9-week old ob/+ or ob/ob mice is shown. C. The average luciferase level relative to body weight of ob/+ vs. ob/ob mice is shown for each of the constructs. A diagram for each sequence is shown on the right. For A-C, WT ob/+ n=9, WT ob/ob n= 4, DR1 ob/+ n=3, DR1 ob/ob n=3, LOF ob/+ n=6, LOF ob/ob n=3, EXT ob/+ n=10, EXT ob/ob n=5.

In contrast, and relative to control mice, the increase in the levels of luciferase expression in nine-week-old ob/ob mice carrying the LOF reporter was 20.6 fold lower (3.7×10^8^ p/s (total flux)) than the increase in nine-week-old ob/ob mice carrying the wild type reporter. Similarly, in ob animals carrying the EXT reporter construct, the increase of luciferase expression in ob/ob mice was 15.8 fold lower (4.8×10^8^ p/s (total flux)) than the increase in luciferase expression in ob/ob mice with the wild type reporter (Fig 5B and Fig 5C).

We found that the increased luciferase expression with obesity was impaired (albeit to a lesser extent) in the DR1 BACTG reporter mice, as the increased expression (3.9×10^9^ p/s (total flux)) of this reporter construct in ob/ob mice was 1.9 fold lower than the expression in ob/ob mice from the wild type construct (Fig 5B and Fig 5C). To reconfirm that the induction of luciferase expression from the LepRE1 DR1 reporter line was impaired, a second cohort of eleven-week-old animals was analyzed. Eleven-week-old ob/ob and ob/+ mice had average body weights of 49g and 25g, respectively. While the body weights of the LepRE1 WT and DR1 ob/ob animals were similar, the increase in luciferase expression from the DR1 reporter (5.3×10^9^ p/s (total flux)) was 1.7 fold lower than the increase in expression from the wild type reporter (9.1×10^9^ p/s (total flux)) (Fig S3).

Finally, we tested whether additional factors in a nuclear extract from ob/ob adipose tissue can interact with the PPARγ/RXRα: LepRE1 complex. We found that addition of a nuclear extract supershifted the PPARγ/RXRα: LepRE1 complex in an EMSA assay while nuclear extracts from wild type adipose tissue did not (see Fig. S4).

In summary, while leptin reporter mice with mutations in LepRE1 show fat-specific expression of luciferase (albeit with lower baseline levels), all three mutants show an impaired response to nutritional changes. The DR1 and EXT mutations fail to reduce luciferase expression after fasting and also show an impairment in the increased expression normally seen on an ob/ob background. The LOF BACTG reporter mice show a lesser reduction of reporter expression after weight loss and also show a profound defect in the increase of luciferase expression with obesity. Overall, these data suggest that trans factors binding to this non-canonical PPARγ binding site and the adjacent EXT sequence play a critical role in controlling the quantitative expression of the leptin gene.

## Discussion

Quantitative control of leptin expression is critical for the homeostatic control of adipose tissue mass. However neither the transcription mechanism nor the signal transduction pathway that regulates the level of leptin expression are known. Their identification has been confounded by the finding that the key regulatory elements regulating leptin expression appear to be responsible for its tissue-specific expression in fat. We thus sought to identify transcription factor binding sites whose footprints (reflecting occupancy) differed in obese vs. wild type animals (Fig 1A). Here we report the identification and functional characterization of a specific PPAR γ binding site (LepRE1) that is responsible for the quantitative control of the leptin gene without affecting its fat-specific expression.

Previous studies from our laboratory have shown that redundant sequences in the extreme 5’ and 3’ regions of the gene, greater than 10kb from the TSS, can confer fat-specific expression of the leptin gene (18, 19). However, because mutations in these regions interfered with the fat-specific expression of this gene, it was impossible to define the sequences that quantitatively regulated leptin expression. This problem also applied to studies of other transcription factors, including C/EBP**α** and SP1 (20, 21), FOSL2 (22) and NFY (18), each of which have been reported to play a role in leptin expression in adipose tissue. Thus, in contrast to the binding sites for these other factors, LepRE1 and the factors that bind to it uncouples the mechanisms conferring quantitative expression of the leptin gene from its fat-specific expression.

LepRE1, and a functionally redundant element in the 3’ region of the leptin gene (LepRE2, Fig 2), show weak binding to PPARγ -- raising the possibility that an additional stabilizing factor is necessary for its binding. PPARγ binding to the LepRE1 and LepRE2 sites are not seen in macrophages (Fig S1)(34) adding further evidence that there is an accessory factor that enables binding in adipocytes. PPARγ, forming an obligate heterodimer with RXRα, is a master regulator of adipogenesis (27, 28). PPARγ expression is highly correlated with leptin expression (38). And an adipose specific PPARγ deletion reduces leptin expression, although this reduction is thought secondary to a defect in lipoatrophy (39). However, PPARγ by itself is not sufficient for a high level of leptin expression. For example, leptin expression is extremely low in cultured adipocytes despite the fact that they express high levels of this transcription factor (16). In addition, thiazolidinediones were found to inhibit leptin expression despite activating PPARγ in cultured adipocytes and rodents(35–37).

The finding of a non-canonical-PPARγ/RXRα-binding sequence (LepRE1) and the effect of cognate mutations in impairing the nutritional regulation of the leptin gene provide evidence that an additional factor(s) is necessary for PPARγ regulated expression. The canonical PPARγ/RXRα binding PPRE motif known as the DR1 sequence is a strong target for binding of this transcription factor (29). The LepRE1 contains an RXRα-binding sequence identical to that in the PPRE (ARE7), but the other half is diverged (Fig 1B). This alteration explains why this sequence was not identified as a binding site by current algorithms. These sequence changes also render the PPARγ/RXRα binding much weaker and explain why PPARγ alone is not sufficient to induce a high level of leptin expression in cultured adipocytes. Furthermore, a PPARγ ligand rosiglitazone (and a RXRα ligand 9-cis-Retinoic acid) decreases the expression of both LE1-driven and LE2-driven luciferase activity (Fig 3A). Such results could explain why thiazolidinediones inhibit leptin expression despite thought as PPARγ agonists. These data are also consistent with previous results with the Estrogen receptor where ligands can reduce expression of a target gene in the absence of a functional co-activator(40). While there were other potential non-canonical PPARγ/RXRα binding sites in the LE1 region, LepRE1 is the only one that was identified in the footprinting studies using ATAC-seq. Nonetheless, even though LepRE1 site was functionally validated and found to be necessary as mentioned above, it is still possible that other sites could also contribute.

A functional requirement of the PPARγ/RXRα complex for quantitative transcriptional regulation of leptin by binding to LepRE1 is suggested by the following evidence: i. Purified PPARγ/RXRα proteins bound IRDye700-labeled LepRE1 with sequence specificity in an (*in vitro*) EMSA assay (Fig 1C, and Fig 1D); ii. ChIP-seq analysis identified PPARγ binding to LepRE1 *in vivo* (31–34)(Fig 1A, and Fig S1B); iii. At baseline, the level of luciferase expression in the LepRE1 LOF reporter mice was considerably lower with a fat-specific luciferase expression level 4~8% that of the wild type construct (Fig 3, and Fig S2); iv. The LOF BACTG reporter mice showed a lesser reduction of reporter expression after fasting and an impairment in the increase of luciferase expression in ob/ob animals (Fig 5).

As mentioned above these data suggest that an additional factor is required to stabilize PPARγ binding to this site to regulate the quantitative level of leptin expression. Our data further suggest that this putative accessory factor binds to the adjacent extension sequence (and potentially part of the nearby DR1 half-site for PPARγ) because mutations in this sequence (LepRE1 EXT) and the nearby PPARγ binding sequence (LepRE1 DR1) do not alter the specificity of reporter expression in fat but do impair the effect of fasting or obesity. The mutation in the extension sequence also dramatically decreases the baseline level of reporter expression, further suggesting that it provides a binding site for a factor that is necessary for high-level expression of leptin in vivo. We also noticed that fat nuclear extracts from ob/ob mice can super shift a purified PPARγ/RXRα: LepRE1 complex in an EMSA assay (Fig S4), suggesting that an additional factor(s) in ob/ob adipocytes is part of the complex. Efforts to identify this other factor(s) are underway. However, we have not ruled out the possibility that a conformational change in PPARγ itself could potentially affect its binding to the extension site as the PPARγ DNA binding domain includes a C terminal helix that inserts into the minor groove of this extension sequence (as shown in the PPARγ/RXRα co-crystal structure)(41). There are numerous other instances in which gene expression is controlled by the stabilization of weak binding. For example, *E. coil* RNA polymerase alone binds fairly weakly to the classic lac promoter and requires the cooperative binding with another low affinity partner (cAMP-CAP) for high level expression of b-galactosidase in the absence of glucose(42). The requirement for accessory factors to facilitate PPARγ-mediated gene expression also has a precedent in brown fat (43). Brown fat expresses high levels of PPARγ but does not express UCP1 unless the PGC-1 coactivator is also expressed though activation of the UCP1 promoter in brown fat also involves other factors such as PRDM16, MED1, and HDAC3 (34, 44, 45). Thus, the stabilization of PPARγ at a non-canonical site may provide a general mechanism for the control of a wide array of other PPARγ target genes. Indeed, previous studies have suggested that the 5’ extension site of PPARγ in the classic DR1 motif may indeed be involved in binding to other factors (30). Structural studies of the glucocorticoid receptor has shown how co-regulatory proteins can alter transcription factor conformation and sequence selection(46). The nature of the accessory factor regulating leptin expression is unknown and under intense investigation. It is noteworthy that while the RXRα-binding sequences are identical between LepRE1 and PPRE (ARE7) of the Fabp4/aP2 gene, LepRE1 has a unique 5’ extension sequence that differs from the extension sequence of the PPARγ site that regulates the UCP1 gene.

We found an impairment in the decrease in reporter expression from the LepRE1 mutants after fasting, as well as a markedly diminished absolute increase in reporter expression from all three of LepRE1 mutants after breeding to ob/ob mice. However, there was still a small relative increase in the expression from the LOF and EXT reporter constructs with obesity (Fig 5C). This suggests that there may exist additional pathways that can partially up-regulate leptin transcription in ob/ob mice. This may be similar to the compensatory increase of leptin transcription from wild type mice to ob/+ mice. Because the body weight difference between wild type and ob/+ mice is almost indistinguishable, this additional LepRE1-independent pathway may not be associated with lipid content in fat.

The most parsimonious model to explain our findings is that the quantitative or qualitative state of an accessory factor(s) that binds to the extension site is altered in concert with changes in fat mass (or something that correlates with these changes) and in turn regulates the binding of PPARγ/RXRα binding to LepRE1 -- thus controlling transcription of the leptin gene (Fig 6). The identification of this factor could thus potentially illuminate the nature of the adipocyte signal transduction pathway that is responsible for the regulated expression of leptin in parallel with changes in cellular lipid content. While several lines of evidence have suggested that cellular lipid content is sensed in adipocytes, the cellular mechanisms are not known. Lipid sensing has also been invoked as potentially regulating the activity of hypothalamic neurons and hepatic metabolism, although the underlying mechanism in these cells types is similarly unknown (47). Possible mechanisms could include regulation of a lipid metabolite, sensing of cell size (which would increase as lipid accumulates), effects of oxygen (the partial pressure of which could vary based on the distance of the nucleus from the capillaries) or other mechanisms. This mystery is analogous to the cholesterol-sensing problem for cells, which was resolved by defining the regulatory mechanisms and signal transduction pathway that regulates the transcription of LDL receptor (11, 12, 48, 49). Identification of the putative accessory factors that bind to the non-canonical PPARγ/RXRα site that we have implicated in the regulation of leptin expression could help resolve this conundrum and lead to the identification of the signal transduction mechanism that links changes in cellular lipid content, or its surrogate, to changes in gene expression and possibly other cellular functions. A deeper understanding of this putative lipid sensing mechanism could be of general importance for understanding the mechanisms responsible for nutritionally mediated changes in cell function and gene expression.

**Fig 6.**
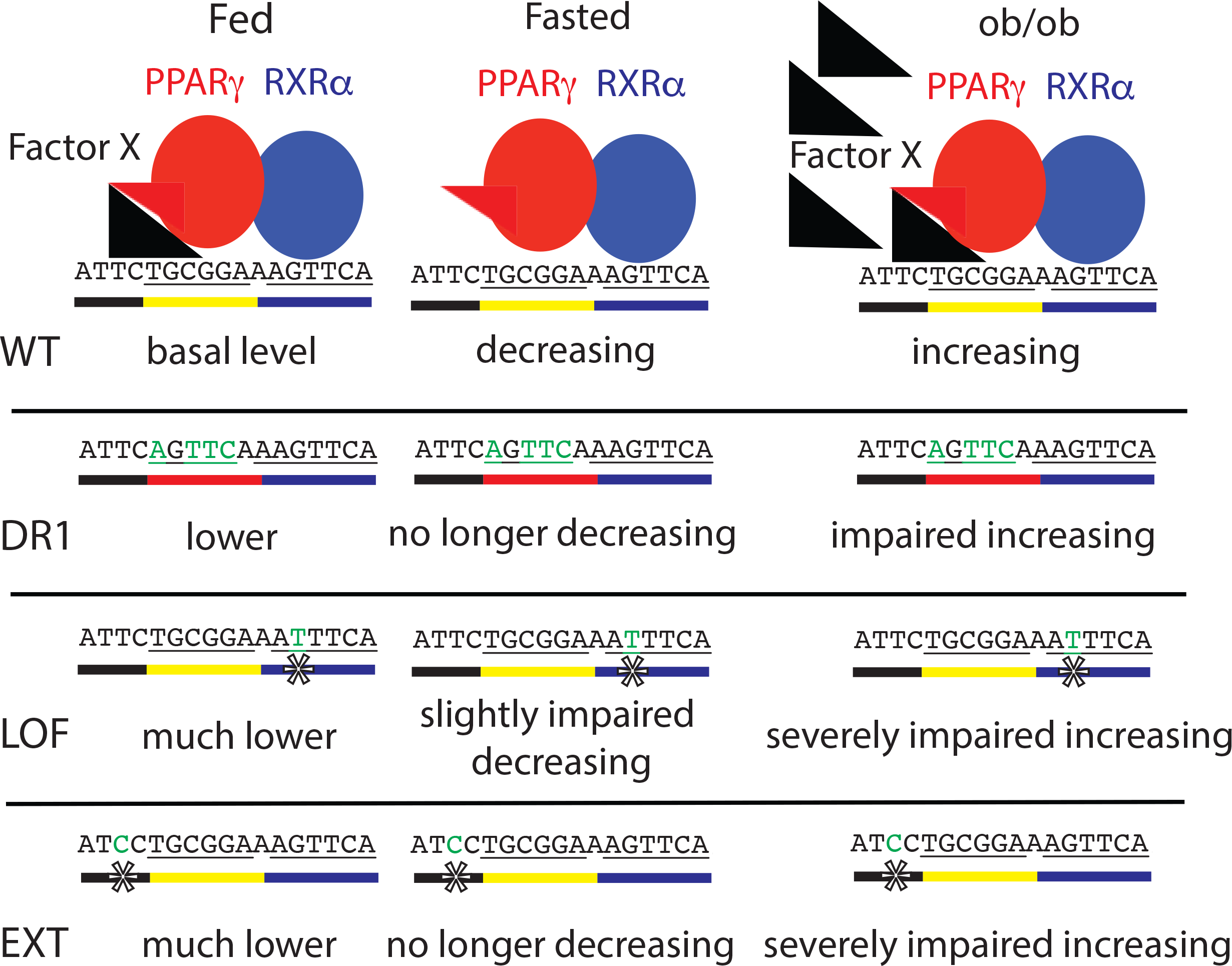
Summary of Leptin Reporter Mice with LepRE1 Mutations and Model for the Transcriptional Regulation of Leptin through a week PPARγ/RXRα binding site LepRE1. The data support a model in which PPARγ/RXRα (red and blue) binding to LepRE1 is stabilized by another factor (black triangle). The mutated nucleotides are green and an asterisk marks the mutations. A red line below the sequence indicates the canonical PPARγ binding site, while a yellow line shows non-canonical PPARγ binding. A blue line below shows RXRα binding sequence. A black line below represents the 5’ extension sequence.

## Materials and Methods

**Experimental animals.** All experiments were approved by The Rockefeller University Institutional Animal Care and Use Committee and were in accordance with the National Institutes of Health guidelines. Both male and female mice (> 3 weeks old) were used for all studies. Mice were housed in a 12 hour light-dark cycle with ad libitum access to food and water, except for fasting assays. All mouse lines are either C57BL/6J or FVB/NJ background. Male mice were used for ATAC-seq studies. Male and female mice were used for luciferase studies.

**ATAC-seq.** 10-week-old C57BL/J6 and B6 ob/ob male mice were purchased from the Jackson Laboratory. Wild type mice were fed or fasted for 2 days. Subcutaneous iWAT was isolated from fasted, fed and ob/ob mice. The fat tissue was minced with blades, dounced in homogenization buffer (20mM Tricine pH7.8, 25mM D-Sucrose, 15mM NaCl, 60mM KCl, 2mM MgCl2, 0.5mM Spermidine), filtered with EMD Millipore Nylon-Net Steriflip™ Vacuum Filter Unit (100um), and centrifuged at 18000g for 5min at 4C. The resulting nuclei pellet was washed once with Tagmentation DNA buffer (10mM Tris pH7.6: acetic acid, 5mM MgCl2, 10% dimethylformamdie)(50) followed by centrifugation at 500g for 5min at 4C. After resuspension in the same Tagmentation DNA buffer and counting using Trypan Blue, an aliquot contacting 50K nuclei was performed transposition reaction using Nextera DNA Library Preparation Kit (illumina) and processed as descripted(23). Library from each mouse was sequenced in one lane using 50bp x2 paired-end reads on an Illumina HiSeq 2500.

Reads were aligned to the mm9 build and Ensemble gene model (NCBIM37) using Bowtie with parameters −X2000 and −m1. Then we processed the alignments using Samtools and adjusted the read start sites to represent the center of the transposon binding event as previously described(23): all reads aligning to the + strand were offset by +4 bp, and all reads aligning to the – strand were offset −5 bp. Reads in two libraries from two individual mouse in each condition were mixed. After that, we found peaks in ob/ob samples with at least a 3 fold higher signal than fasted conditions using Homer (51). Within the above peak region, footprint scores were calculated based Wellington method(52). The conservation score within mammals was downloaded from a UCSC server and calculated from 20 placental mammal species(53). All the results were viewed using the Integrative Genomics Viewer (IGV)(54).

**Purification of PPARγ and RXRα**. Baculovirus expressing recombinant FLAG-tagged mouse PPARγ2 (F-mPPARγ2) and human RXRα (F-hRXRα) were prepared as described(55). Sf9 cells were infected with each baculovirus to express the recombinant protein separately and the soluble extract was prepared by sonication in BC-buffer (20 mM Hepes-KOH (pH7.9), 1 mM EDTA, 10% glycerol, 1 mM DTT, and 0.5 mM PMSF) containing 100mM KCl. Clear lysate after the ultra centrifugation was subjected to HiTrap-Q (GE healthcare) and bound proteins were eluted by liner gradient of KCl. PPARγ2 and RXRα were eluted from 250 mM to 300 mM KCl, and from 150 mM to 200 mM KCl, respectively. PPARγ2 or RXRα were further purified by M2-agarose (Sigma) in BC-buffer containing 300mM KCl, 0.1 % NP40, and 0.25 mM DTT, and eluted by 0.15mg/ml triple FLAG peptide (Sigma).

**EMSA.** EMSA was performed using the LICOR Odyssey EMSA Buffer Kit. Basically, purified PPARγ (10ng) and RXRα (10ng) were incubated in a 20ul reaction volume with 5mM Tris pH 7.5, 25mM KCl, 3mM DTT, 0.25% Tween 20, 5mM MgCl2, 1ug Poly (dI’dC) (or 0.5ug Salmon Sperm DNA), 2.5nM IRDye 700 labeled DNA probe, with or without 500nM unlabeled oligos for 20 min at room temperature. Samples were then mixed with 2ul 10X LICOR orange loading dye and loaded onto a 5% Mini-PROTEAN^®^ TBE Gel (BioRad). After ~75min run at 70V, the gel was scanned with LICOR Odyssey CLx Imaging System. The results were analyzed and quantified with Image Studio Lite (LICOR). The sequence of the IRDye 700 labeled and unlabeled oligos are shown in Table S1, respectively. All oligos in this paper were ordered from Integrated DNA Technologies, Inc.

**Plasmids.** pTK-Renilla luciferase encoded the Renilla luciferase gene under control of the TK promoter (Promega). pCMV-PPARγ encoded the mouse PPARγ2 under control of the CMV promoter(55). pCMV-RXRα encoded FLAG-tagged human RXRα under control of the CMV promoter(55). pLE1-firefly luciferase was generated by cloning 115bp leptin enhancer 1(LE1)(GAGAACACTTAACAGCAAAGGTTAATCTTTGAAGTCCCTAAAGATTTGAACTTTCCGCAGAATTGGCTGCAGCGTCTAGTGGGTTAGAGTCTAATTGGAGTAGAGCAGAAGCAAG) into pGL4.27 (Promega) between XhoI and HindIII sites. pLE2-firefly luciferase was generated by cloning 279bp leptin enhancer 2 (LE2)(TGGAGGGGCTTTTGGAGAGCTGTTTGTGTGTGACAGGGCAAGGCCTGGCTGGCGTCCAGCCATCACCAGGGTCAGCCCCACCCGGGCTTGGCCACAGCCAGCTACCAGTTATTCAGGCGAAAGGATTACCCTAAGCCCAGGGCCAGGCAAGAAGCAAATTCTACACCAGCGGCTGAGCAGTTCTGCAAACCAGCCTCGAGAAGCACCCAGTTATTTTTAAAGCCAGAGTATCAAAACCCCAAGCAAATAACCAAACCCAAACCTCACAGTCTAATGGCA) into pGL4.27 (Promega) between BgIII and HindIII sites.

**Dual Luciferase Reporter Assay.** The reporter assay was performed as described (56, 57). On day 0, HEK293T cells were set up for experiments in 0.5ml of DMEM (Corning, 10-013-CV) with supplemented with 10% FBS (Thermo Fisher Scientific, 16000044) at a density of 30,000 cells/well in 24-well plates. On day 1, cells were co-transfected with 0.125ng of pTK-Renilla luciferase, 12.5ng pLE1-firefly luciferase or pLE2-firefly luciferase, 162.7ng of pCMV-PPARγ, 162.7ng of pCMV-RXRα. Fugene 6 was used as the transfection agent. For each transfection, the total amount of DNA was adjusted to 338ng/dish by the addition of pcDNA mock vector. On day 2, the cells were treated with 1uM rosiglitazone (Sigma) (in DMSO), 1uM 9-cis-Retinoic acid (Sigma) (in DMSO), or DMSO alone. On day 3, the cells were washed with PBS, after which luciferase activity was read on a CLARIOstar (BMG Labtech) using the Dual-Luciferase Reporter Assay System (Promega). The amount of LE1 (or LE2) luciferase activity in each dish was normalized to the amount of Renilla luciferase activity in the same dish. Relative luciferase activity of 1 represents the normalized luciferase value in dishes transfected with pcDNA mock vector with DMSO treatment. All values are the average of duplicate assays.

**BAC modification.** Recombineering was performed as previously described (58) on a BAC containing −22 kb to +8.8 kb leptin-luciferase reporter construct (18). Sequences of primers for creating mutations are included in Table S1. Genomic sequence and coordinates were based on NCBI37/mm9 mouse genome.

**Transgenic animals.** Leptin-luciferase reporter BACs were used to generate transgenic animals in the inbred FVB N/J background (Jackson Lab) using common pronuclear injection techniques (17, 59). B6 ob/+ mice (Jackson Lab) mice were crossed to FVB N/J mice (Jackson Lab) for many generations to generate fully inbred FVB ob/+ mice. Leptin-luciferase reporter BAC transgenic animals were mated with above FVB ob/+ mice to produce FVB ob/+ and FVB ob/ob transgenic animals.

***In vivo* luciferase imaging.** 50ul 15mg/ml of XenoLight D-Luciferin (PerkinElmer) in 1xDPBS was injected intraperitoneally into awake mice. After mice were anesthetized with isoflurane, sequential images were taken with IVIS Spectrum In Vivo Imaging System (PerkinElmer) until the luciferase intensity passed the highest point. Imaging was normally performed within 15min after luciferin injection. The photon image was later analyzed by Living Image 4.5 software (PerkinElmer). The image of the highest luciferase intensity was chosen for further analysis.

***In vitro* luciferase assay.** Tissues were harvested and homogenized using POLYTRON^®^ PT 1200E Handheld Homogenizer. After centrifugation at 20000g for 10min at 4C, the supernatant was loaded to a 96 well plate and measured using the CLARIOstar(BMG Labtech). We used Luciferase Reporter Assay System (Promega) for luciferase assay and DC™ Protein Assay (BioRad) for protein concentration measurement.

**Quantification and Statistical analysis.** Mean values are accompanied by SEM. Statistical parameters including the sample size (n = number of animals or samples per group), precision measures (mean ± SEM) and statistical significance are reported in the Figs and Fig Legends. Two-ended, unpaired Student T test was used. Significance was defined as p < 0.05. Significance annotations are: *p < 0.05, **p < 0.01, ***p < 0.001, ****p < 0.0001. Mice were randomized into control or treatment groups. Control mice were age-matched littermate controls where possible. All statistics and data analysis were performed using GraphPad Prism 7.

**Data Resources.** The ATAC-seq data generated in this publication can be found online associated with GEO Publication Reference ID (GSE113413).

**AUTHOR CONTRIBUTIONS**. Y.Z. conceived/designed the study, performed the majority of the experiments, and analyzed all the data. O.S.D developed the strategy and reagents for generating BAC Transgenic Mice. T.N. performed the purification of PPARγ and RXRα proteins. G.F. performed pronuclear injections for BACTGs and some of the in vivo luciferase imaging. Y.L. provided some experimental advice and help. M.A.L. helped the PPARγ ChIP-seq analysis. R.G.R. conceived part of the study. J.M.F. conceived the majority of the study. Y.Z. and J.M.F. wrote the manuscript with input from all authors.

## ACKNOWLEDGEMENTS

We thank Kivanc Birsoy, Paul Cohen, Henrik Molina, Sohail Malik, and Ming Yu for valuable discussions, Kenneth Lay and Elaine Fuchs for help with LICOR Odyssey CLx imaging system, Wenxiang Hu (U. Pennsylvania) and Chunjie Jiang (U. Pennsylvania) for bioinformatic discussions, Xiaofei Yu for help with cell culture, the Rockefeller University Genomics Resource Center and Comparative Bioscience Center. This project was supported by funding from the JPB Foundation (CEN 5402133; J.M.F.) and NIH (R01-DK071900; R.G.R.). Y.Z. acknowledges support from the Howard Hughes Medical Institute. O.S.D. acknowledges support from the Swedish Research Council Fellowship and The Swedish Medical Research Society Fellowship. M.A.L. acknowledges support from NIH (R01-DK049780) and the JPB Foundation.

